# Vein Patterning by Tissue-Specific Auxin Transport

**DOI:** 10.1101/866632

**Authors:** Priyanka Govindaraju, Carla Verna, Tongbo Zhu, Enrico Scarpella

## Abstract

Unlike in animals, in plants vein patterning does not rely on direct cell-cell interaction and cell migration; instead, it depends on the transport of the plant signal auxin, which in turn depends on the activity of the PIN-FORMED1 (PIN1) auxin transporter. The current hypotheses of vein patterning by auxin transport propose that in the epidermis of the developing leaf PIN1-mediated auxin transport converges to peaks of auxin level. From those convergence points of epidermal PIN1 polarity, auxin would be transported in the inner tissues where it would give rise to major veins. Here we tested predictions of this hypothesis and found them unsupported: epidermal PIN1 expression is neither required nor sufficient for auxin-transport-dependent vein patterning, whereas inner-tissue PIN1 expression turns out to be both required and sufficient for auxin-transport-dependent vein patterning. Our results refute all vein patterning hypotheses based on auxin transport from the epidermis and suggest alternatives for future tests.

## Introduction

Most multicellular organisms solve the problem of long-distant transport of signals and nutrients by means of tissue networks such as the vascular system of vertebrate embryos and the vein networks of plant leaves; therefore, how vascular networks form is a key question in biology. In vertebrates, the formation of the embryonic vascular system relies on direct cell-cell interaction and at least in part on cell migration (e.g., (Noden, 1988; Xue et al., 1999)), both of which are precluded in plants by a wall that keeps cells apart and in place; therefore, vascular networks form differently in plant leaves.

How leaf vein networks form is unclear, but available evidence suggests that polar transport of the plant signal auxin is non-redundantly required for vein patterning (Mattsson, Sung, & Berleth, 1999; Sieburth, 1999). Such non-redundant functions of polar auxin transport in vein patterning in turn depend on non-redundant functions of the PIN-FORMED1 (PIN1) auxin transporter (Galweiler et al., 1998; Petrasek et al., 2006; Zourelidou et al., 2014; Sawchuk, Edgar, & Scarpella, 2013; Verna, Ravichandran, Sawchuk, Linh, & Scarpella, 2019). At early stages of leaf development, PIN1 polar localization at the plasma membrane of epidermal cells is directed toward single cells along the marginal epidermis (Reinhardt et al., 2003; Benkova et al., 2003; Heisler et al., 2005; Scarpella, Marcos, Friml, & Berleth, 2006; Wenzel, Schuetz, Yu, & Mattsson, 2007; Hay, Barkoulas, & Tsiantis, 2006; Bayer et al., 2009). These convergence points of epidermal PIN1 polarity are associated with broad domains of PIN1 expression in the inner tissue of the developing leaf, and these broad domains will over time become restricted to the narrow sites where the midvein and lateral veins will form.

Consistent with those observations, the prevailing hypotheses of vein patterning propose that convergence points of epidermal PIN1 polarity contribute to the formation of local peaks of auxin level in the epidermis, and that that auxin is transported by PIN1 from the epidermal convergence points into the inner tissues of the leaf, where it will lead to vein formation (reviewed in (Runions, Smith, & Prusinkiewicz, 2014; Prusinkiewicz & Runions, 2012); see also (Alim & Frey, 2010; Hartmann, Barbier de Reuille, & Kuhlemeier, 2019), and references therein). As such, these hypotheses predict that epidermal PIN1 expression is required for vein patterning. Here we tested this prediction and found it unsupported: epidermal PIN1 expression is neither required nor sufficient for auxin-transport-dependent vein patterning; instead, PIN1 expression in the inner tissues turns out to be both required and sufficient for auxin-transport-dependent vein patterning. Our results refute all the current hypotheses of vein formation that depend on polar auxin transport from the epidermis and suggest alternatives for future testing.

## Results and Discussion

### PIN1 Expression during Arabidopsis Vein Patterning

In Arabidopsis leaf development, the formation of the midvein precedes the formation of the first loops of veins (“first loops”), which in turn precedes the formation of the second loops (Mattsson et al., 1999; Sieburth, 1999; Sawchuk, Head, Donner, & Scarpella, 2007; Scarpella, Francis, & Berleth, 2004; Kang & Dengler, 2004) (Fig. 1A–C). The formation of second loops precedes the formation of third loops and that of minor veins in the area delimited by the midvein and the first loops (Fig. 1C,D). Loops and minor veins form first near the top of the leaf and then progressively closer to its bottom, and minor veins form after loops in the same area of the leaf (Fig. 1B–D).

**Figure 1.**
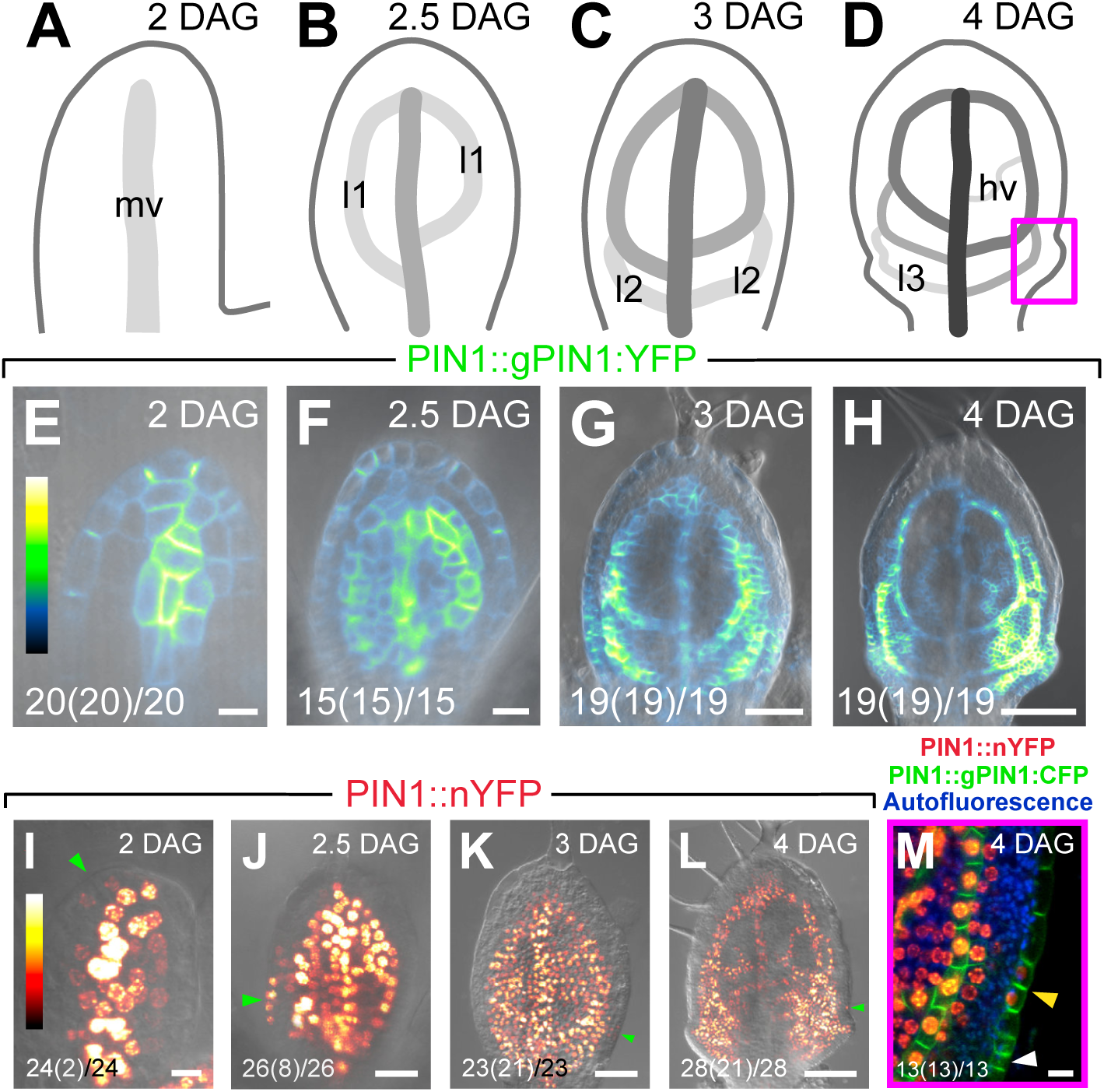
PIN1 Expression During Arabidopsis Vein Patterning. (A–M). Top right: leaf age in days after germination (DAG). Abaxial side to the left in (A,E,I). (A–D) Midvein, loops, and minor veins form sequentially during leaf development (Mattsson et al., 1999; Sieburth, 1999; Sawchuk et al., 2007; Scarpella et al., 2004; Kang & Dengler, 2004); increasingly darker grays depict progressively later stages of vein development. Box in (D) illustrates position of closeup in (M) and in Figs. 2D,J and 4D,J. (E–M) Confocal laser scanning microscopy with (E–L) or without (M) transmitted light. Bottom left: reproducibility index, i.e. no. of leaves with the displayed inner-tissue expression (no. of leaves with the displayed epidermal expression) / no. of leaves analyzed. Lookup tables in (E–H) — ramp in (E) — and in (I–L) — ramp in (I) — visualize expression levels. Green arrowheads in (I–L) and yellow arrowhead in (M) point to epidermal expression; white arrowhead in (M) points to convergence point of PIN1 polarity. hv, minor vein; l1, first loop; l2, second loop; l3, third loop; mv, midvein. Scale bars: (E,I,M) 10 μm; (F,J) 20 μm; (G,K) 50 μm; (H,L) 100 μm.

Consistent with previous reports (Scarpella et al., 2006; Wenzel et al., 2007; Sawchuk et al., 2013; Heisler et al., 2005; Benkova et al., 2003; Marcos & Berleth, 2014; Bayer et al., 2009; Reinhardt et al., 2003; Sawchuk et al., 2007; Verna et al., 2019), a fusion of the *PIN-FORMED1* (*PIN1*) open reading frame to YFP driven by the *PIN1* promoter (PIN1::gPIN1:YFP) (Xu et al., 2006) was expressed in all the cells of the leaf at early stages of tissue development; over time, however, epidermal expression became restricted to the basalmost cells, and inner-tissue expression became restricted to developing veins (Fig. 1E–H).

We asked whether PIN1::gPIN1:YFP expression were recapitulated by the activity of the *PIN1* promoter. To address this question, we imaged expression of a nuclear YFP driven by the *PIN1* promoter (PIN1::nYFP) in first leaves 2, 2.5, 3, and 4 days after germination (DAG).

Just like PIN1::gPIN1:YFP (Fig. 1E–H), PIN1::nYFP was expressed in all the inner cells of the leaf at early stages of tissue development, and over time this inner-tissue expression became restricted to developing veins (Fig. 1I–L). However, unlike PIN1::gPIN1:YFP and PIN1::gPIN1:CFP (Gordon et al., 2007) (Fig. 1E– H,M), PIN1::nYFP was expressed in very few epidermal cells at the tip of 2-DAG primordia and at the margin of 2.5-DAG primordia, and this epidermal expression was very rare (Fig. 1I,J). PIN1::nYFP expression in epidermal cells at the leaf margin was more frequent at 3 and 4 DAG but was still limited to very few cells (Fig. 1K–M). Moreover, these PIN1::nYFP-expressing epidermal cells were not those that contributed to convergence points of epidermal PIN1 polarity (Fig. 1M).

Because a fusion of the *PIN1* coding sequence to GFP driven by the *PIN1* promoter (PIN1::cPIN1:GFP) was hardly expressed in leaf epidermal cells (Fig. 2C,D,I,J), we conclude that the already limited activity of the *PIN1* promoter in the leaf epidermis is suppressed by the *PIN1* coding sequence and that the leaf epidermal expression characteristic of PIN1 is encoded in the gene’s introns.

**Figure 2.**
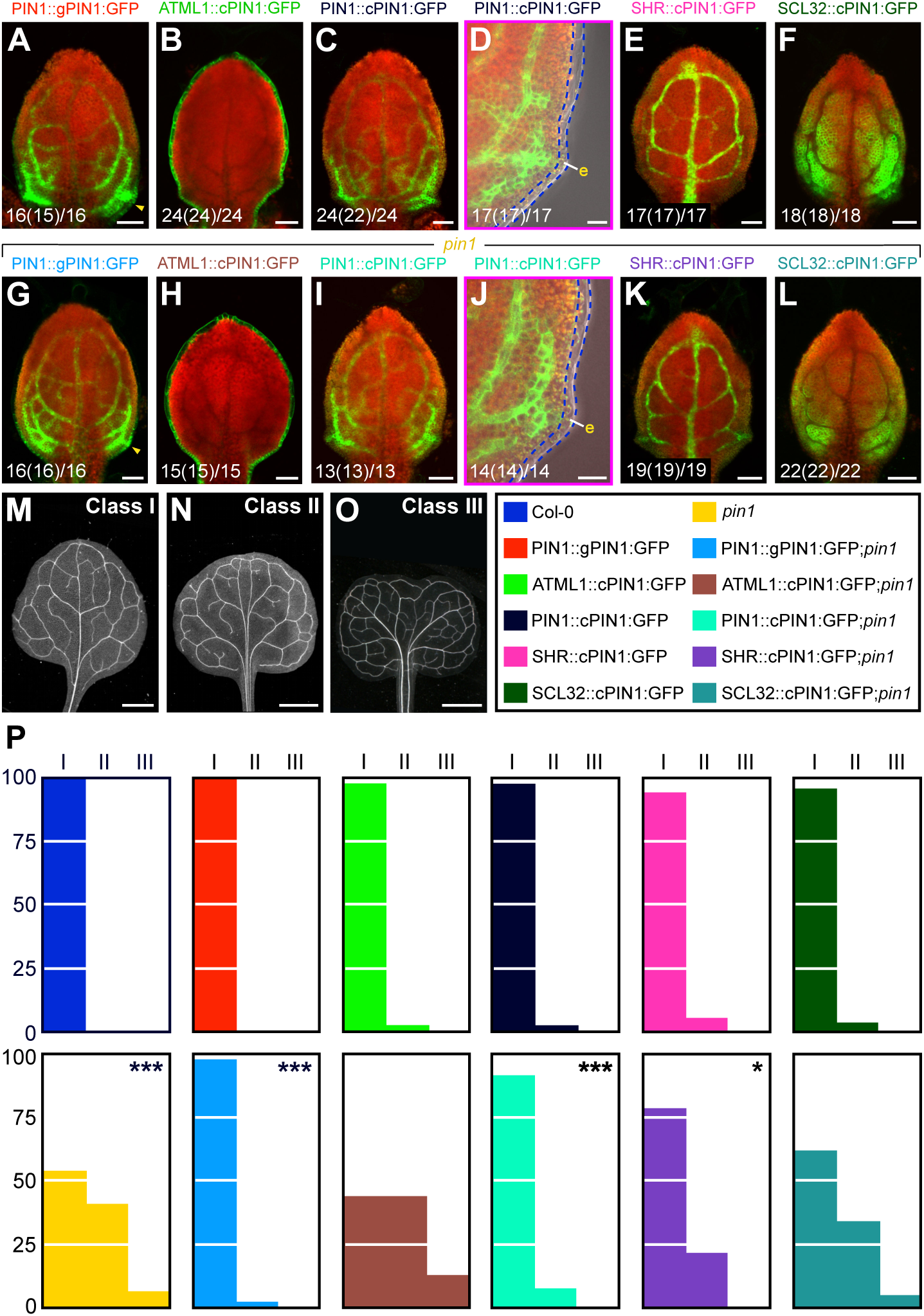
Tissue-Specific PIN1 Expression in *PIN1*-dependent Vein Patterning. (A–L). Confocal laser scanning microscopy with (D,J) or without (A–C,E–I,K,L) transmitted light; first leaves 4 DAG. Green, GFP expression; red, autofluorescence. Yellow arrowheads in (A,G) point to epidermal expression. Bottom left: reproducibility index, i.e. no. of leaves with the displayed inner-tissue expression (no. of leaves with the displayed epidermal expression) / no. of leaves analyzed. (M–O) Dark-field illumination of mature first leaves illustrating phenotype classes (top right): class I, I-shaped midvein (M); class II, Y-shaped midvein (N); class III, fused leaves (O). (P) Percentages of leaves in phenotype classes. Difference between *pin1* and WT, between PIN1::gPIN1:GFP;*pin1* and *pin1*, and between PIN1::cPIN1:GFP;*pin1* and *pin1* was significant at *P*<0.001 (***) by Kruskal-Wallis and Mann-Whitney test with Bonferroni correction. Difference between SHR::cPIN1:GFP;*pin1* and WT, and between SHR::cPIN1:GFP;*pin1* and *pin1* was significant at *P*<0.05 (*) by Kruskal-Wallis and Mann-Whitney test with Bonferroni correction. Sample population sizes: WT, 40; *pin1*, 60; PIN1::gPIN1:GFP, 55; ATML1::cPIN1:GFP, 49; PIN1::cPIN1:GFP, 48; SHR::cPIN1:GFP, 59; SCL32::cPIN1:GFP, 60; PIN1::gPIN1:GFP;*pin1*, 60; ATML1::cPIN1:GFP;*pin1*, 55; PIN1::cPIN1:GFP;*pin1*, 51; SHR::cPIN1:GFP;*pin1*, 60; SCL32::cPIN1:GFP;*pin1*, 58. e, epidermis. Scale bars: (A–C,E–I,K,L) 60 μm; (D,J) 20 μm; (M) 1 mm; (N,O) 2 mm.

### Tissue-Specific PIN1 Expression in *PIN1* Non-Redundant Functions in Vein Patterning

During leaf development, PIN1 is expressed in all the tissues — the epidermis, the vascular tissue, and the nonvascular inner tissue (Figure 1). We asked what the function in *PIN1*-dependent vein patterning were of PIN1 expression in these tissues. To address this question, we expressed in the WT and *pin1* mutant backgrounds

1. PIN1::gPIN1:GFP, which like PIN1::gPIN1:YFP and PIN1::gPIN1:CFP (Fig. 1E–H,M) is expressed in all the tissues of the developing leaf (Fig. 2A,G);
2. cPIN1:GFP driven by the epidermis-specific *ARABIDOPSIS THALIANA MERISTEM LAYER1* promoter (Sessions, Weigel, & Yanofsky, 1999) (ATML1::cPIN1:GFP) (Fig. 2B,H);
3. PIN1::cPIN1:GFP, which is expressed in the leaf inner tissues (Fig. 2C,D,I,J);
4. cPIN1:GFP driven by the vascular-tissue-specific *SHORT-ROOT* promoter (Gardiner, Donner, & Scarpella, 2011) (SHR::cPIN1:GFP) (Fig. 2E,K);
5. cPIN1:GFP driven by the *SCARECROW-LIKE32* promoter, which is active in the nonvascular inner tissue of the leaf (Gardiner et al., 2011) (SCL32::cPIN1:GFP) (Fig. 2F,L).

We then compared vein patterns of mature first leaves of the resulting backgrounds.

Consistent with previous reports (Sawchuk et al., 2013; Verna et al., 2019), the vein patterns of nearly 50% of *pin1* leaves were abnormal (Fig. 2M–P). The vein patterns of PIN1::gPIN1:GFP, ATML1::cPIN1:GFP, PIN1::cPIN1:GFP, SHR::cPIN1:GFP, and SCL32::cPIN1:GFP were no different from the WT vein pattern (Fig. 2M–P). Both PIN1::gPIN1:GFP and PIN1::cPIN1:GFP normalized the phenotype spectrum of *pin1* vein patterns (Fig. 2M–P; Fig. S1A,C). SHR::cPIN1:GFP shifted the phenotype spectrum of *pin1* vein patterns toward the WT vein pattern (Fig. 2M–P; Fig. S1D). The vein pattern defects of ATML1::cPIN1:GFP;*pin1* and SCL32::cPIN1:GFP;*pin1* were no different from those of *pin1* (Fig. 2M–P; Fig. S1B,E). We observed a similar effect of tissue-specific PIN1 expression in *PIN1*-dependent cotyledon patterning (Figure S2).

Consistent with interpretation of similar findings in other organisms (e.g., (Wisidagama, Thomas, Lam, & Thummel, 2019; Cherbas, Hu, Zhimulev, Belyaeva, & Cherbas, 2003; Topalidou & Miller, 2017; Soloviev, Gallagher, Marnef, & Kuwabara, 2011)), we conclude that PIN1 expression in the epidermis is neither required nor sufficient for *PIN1*-dependent vein patterning. By contrast, PIN1 expression in the inner tissues of the leaf is both required and sufficient for *PIN1*-dependent vein patterning; such function of PIN1 expression seems to mainly depend on PIN1 expression in the vascular tissue.

### Expression of PIN3, PIN4, and PIN7 During Vein Patterning

Collectively, *PIN3, PIN4*, and *PIN7* act redundantly with *PIN1* in *PIN1-*dependent vein patterning, and like *PIN1* they are expressed in both epidermis and inner tissues of young leaves (Verna et al., 2019). In those leaves, however, the most reproducible features of the Arabidopsis vein pattern can already be recognized (Donner, Sherr, & Scarpella, 2009; Gardiner, Sherr, & Scarpella, 2010; Gardiner et al., 2011; Sawchuk et al., 2013; Donner & Scarpella, 2013; Verna, Sawchuk, Linh, & Scarpella, 2015; Amalraj et al., 2019; Verna et al., 2019). Therefore, to test the possibility that compensatory functions provided by *PIN3, PIN4*, and *PIN7* may account for the observation that PIN1 expression in the epidermis is dispensable and that PIN1 expression in the inner tissues of the leaf is sufficient for *PIN1*-dependent vein patterning, we first asked what the expression were of PIN3, PIN4, and PIN7 during vein patterning. To address this question we imaged expression of PIN3::gPIN3:YFP, PIN4::gPIN4:YFP, and PIN7::gPIN7:YFP in first leaves 2, 2.5, 3, and 4 DAG.

#### PIN3 Expression

At 2 DAG, PIN3::gPIN3:YFP was expressed in the abaxial epidermis, though more strongly near its top, and in inner cells on the abaxial side of the primordium, mainly at its bottom (Fig. 3A). At 2.5 DAG, PIN3::gPIN3:YFP was expressed in the marginal epidermis, though more strongly near its top (Fig. 3B). Inner expression was restricted to the top and bottom of the midvein and to and around the top of the first loops. At 3 DAG, PIN3::gPIN3:YFP expression persisted in the marginal epidermis, but strong expression had spread toward the bottom of the primordium (Fig. 3C). Inner expression had spread to the whole midvein but was stronger at its top and bottom; inner expression had also spread toward the bottom of the primordium but was stronger in and around the first loops. At 4 DAG, PIN3::gPIN3:YFP expression continued to persist in the marginal epidermis, but strong expression had spread to the whole lamina (Fig. 3D). Inner expression persisted in the midvein and remained stronger at its top and bottom; furthermore, inner expression had spread to the entire lamina but was stronger in and around loops and minor veins.

**Figure 3.**
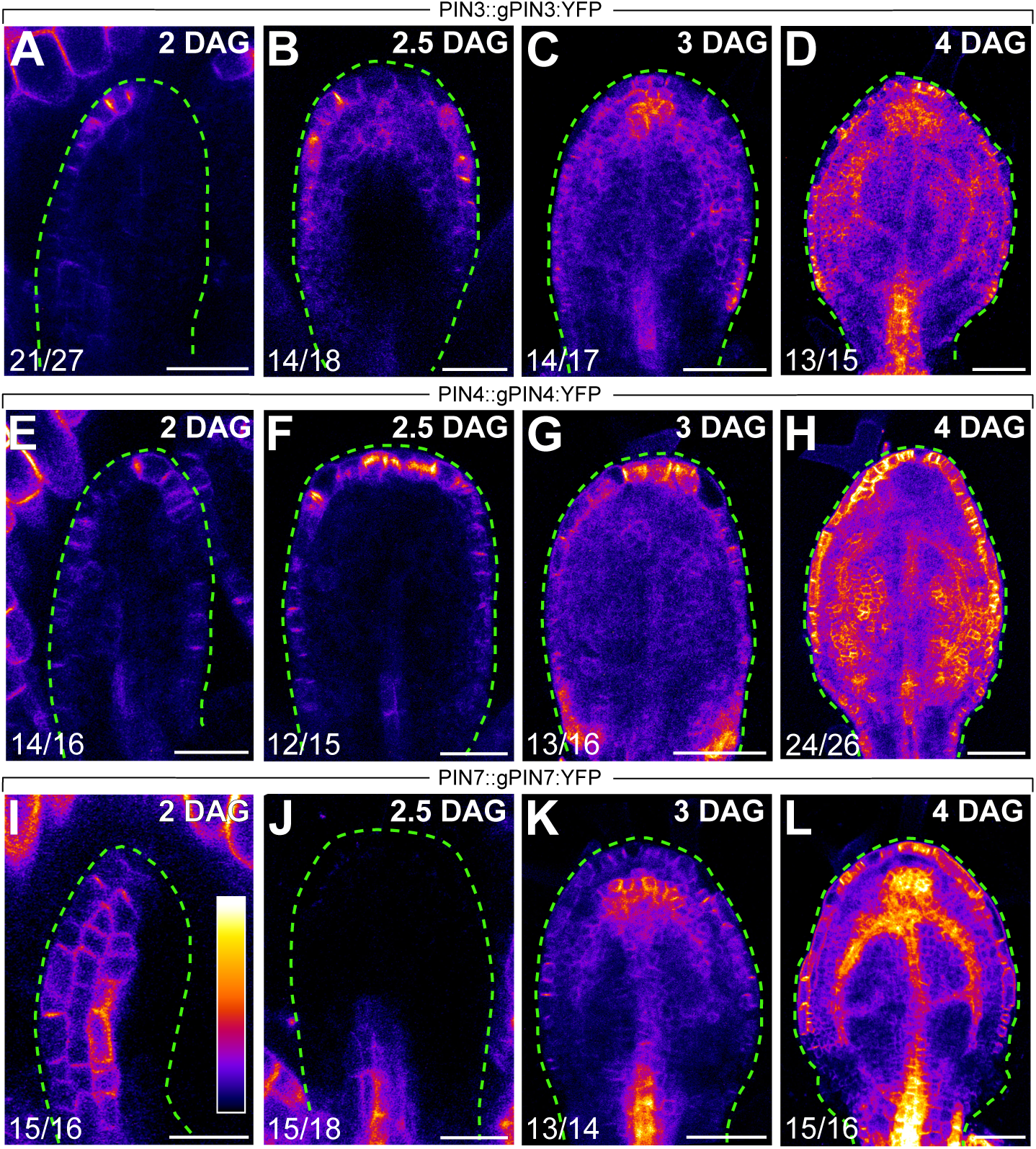
Expression of PIN3, PIN4, and PIN7 During Vein Patterning. (A–L) Confocal laser scanning microscopy. Top right: leaf age in DAG; bottom left: reproducibility index, i.e. no. of leaves with the displayed expression / no. of leaves analyzed. Lookup table — ramp in (I) — visualizes expression levels. Abaxial side to the left in (A,E,I). Scale bars: (A,B,E,F,I,J) 30 μm; (C,D,G,H,K,L) 60 μm.

#### PIN4 Expression

At 2 DAG, PIN4::gPIN4:YFP was expressed in both the adaxial and abaxial epidermis, though more strongly at the top of the primordium (Fig. 3E). Inner expression was restricted to the bottom of the midvein and to very few cells scattered across the primordium. At 2.5 DAG, PIN4::gPIN4:YFP was expressed in the marginal epidermis, though more strongly at its top (Fig. 3F). Inner expression persisted at the bottom of the midvein and in very few cells scattered across the primordium. At 3 DAG, PIN4::gPIN4:YFP expression persisted in the marginal epidermis, though expression was stronger at its top and bottom (Fig. 3G). Inner expression had spread to the whole midvein and to small groups of cells scattered across the primordium. At 4 DAG, PIN4::gPIN4:YFP continued to be expressed in the marginal epidermis, but expression had become more homogeneous (Fig. 3H). Inner expression persisted in the midvein and had spread to and around loops and larger groups of cells scattered across the lamina.

#### PIN7 Expression

At 2 DAG, PIN7::gPIN7:YFP was expressed in the abaxial epidermis and in inner cells on the abaxial side of the primordium (Fig. 3I). At 2.5 DAG, PIN7::gPIN7:YFP was expressed at the bottom of the midvein (Fig. 3J). At 3 DAG, PIN7::gPIN7:YFP became expressed in the marginal epidermis, though expression was stronger near the top of the primordium (Fig. 3K). Inner expression had spread to the whole midvein but was stronger at its top and bottom; inner expression had also spread to and around the first loops, though expression was stronger at their top. At 4 DAG, PIN7::gPIN7:YFP expression had spread to the whole marginal epidermis but was weaker at its bottom (Fig. 3L). Inner expression persisted in the midvein and remained stronger at its top and bottom; furthermore, inner expression had spread to the whole lamina, though expression was stronger in and around loops and minor veins.

In conclusion, during vein patterning PIN3, PIN4, and PIN7 are collectively expressed in the epidermis, in developing veins, and — more weakly — in the nonvascular inner tissue of the leaf.

### Tissue-Specific PIN1 Expression in *PIN1* Redundant Functions in Vein Patterning

Collectively, *PIN3, PIN4*, and *PIN7* act redundantly with *PIN1* in *PIN1-*dependent vein patterning (Verna et al., 2019), and they are expressed in the leaf epidermis and inner tissues during vein patterning (Figure 3). Therefore, to test the possibility that compensatory functions provided by *PIN3, PIN4*, and *PIN7* may account for the observation that PIN1 expression in the epidermis is dispensable and that PIN1 expression in the inner tissues of the leaf is sufficient for *PIN1*-dependent vein patterning, we next expressed in the *pin3;pin4;pin7* (*pin3;4;7* hereafter) and *pin1,3;4;7* mutant backgrounds

1. PIN1::gPIN1:GFP, which is expressed in all the tissues of the developing leaf (Fig. 4A,G);
2. ATML1::cPIN1:GFP, which is only expressed in the epidermis (Fig. 4B,H);
3. PIN1::cPIN1:GFP, which is expressed in the leaf inner tissues (Fig. 4C,D,I,J);
4. SHR::cPIN1:GFP, which is only expressed in the vascular tissue (Fig. 4E,K);
5. SCL32::cPIN1:GFP, which is expressed in the nonvascular inner tissue of the leaf (Fig. 4F,L).

We then compared vein patterns of mature first leaves of the resulting backgrounds.

**Figure 4.**
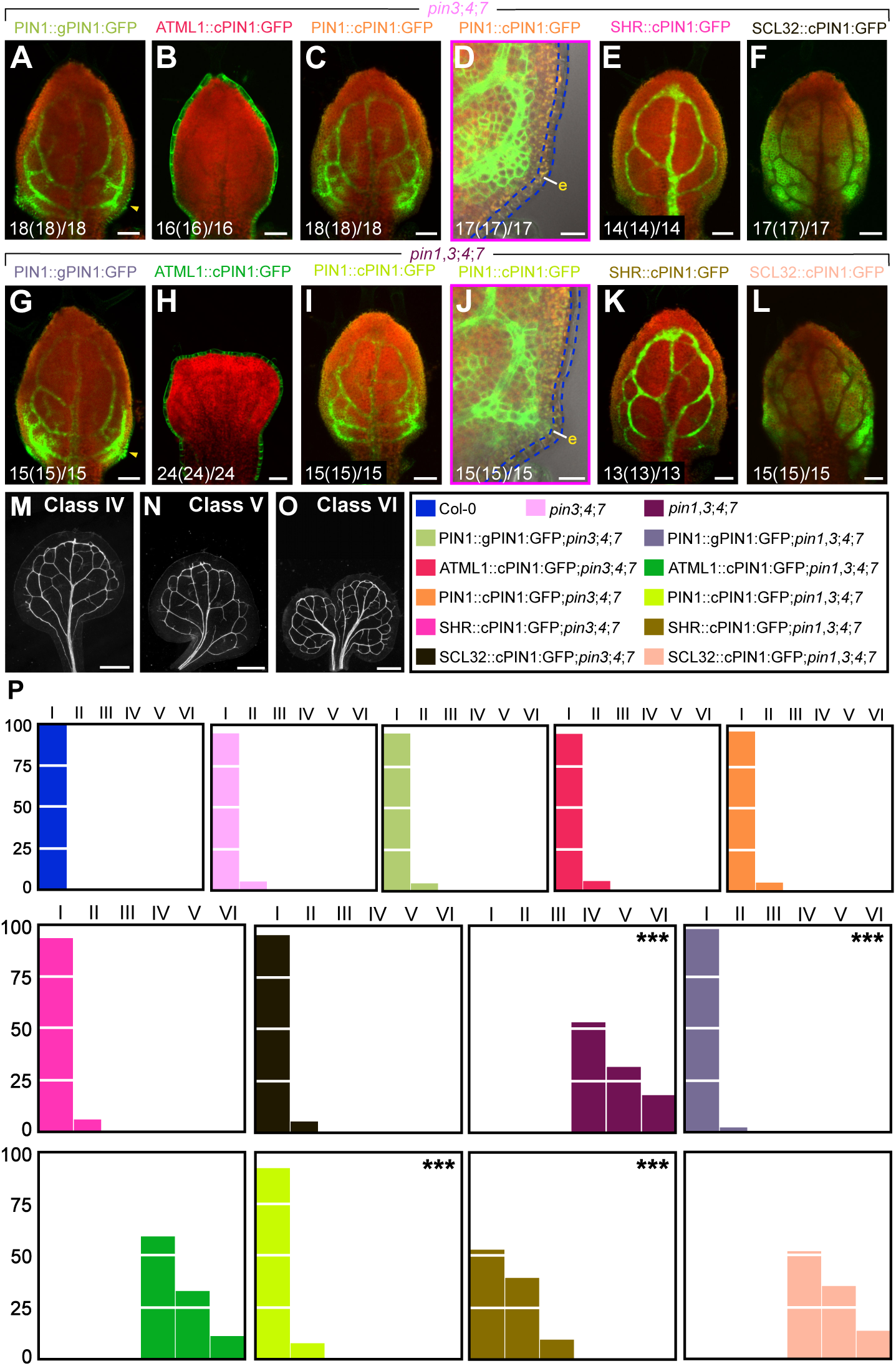
Tissue-Specific PIN1 Expression in *PIN1/PIN3/PIN4/PIN7*-dependent Vein Patterning. (A–L). Confocal laser scanning microscopy with (D,J) or without (A–C,E–I,K,L) transmitted light; first leaves 4 DAG. Green, GFP expression; red, autofluorescence. Yellow arrowheads in (A,G) point to epidermal expression. Bottom left: reproducibility index, i.e. no. of leaves with the displayed inner-tissue expression (no. of leaves with the displayed epidermal expression) / no. of leaves analyzed. (M–O) Dark-field illumination of mature first leaves illustrating phenotype classes (top right): class IV, I-shaped midvein and thick veins (M); class V, Y-shaped midvein and thick veins (N); class VI, fused leaves with thick veins (O). (P) Percentages of leaves in phenotype classes. Difference between *pin1,3;4;7* and WT, between PIN1::gPIN1:GFP;*pin1,3;4;7* and *pin1,3;4;7*, between PIN1::cPIN1:GFP;*pin1,3;4;7* and *pin1,3;4;7*, between SHR::cPIN1:GFP;*pin1,3;4;7* and *pin3;4;7*, and between SHR::cPIN1:GFP;*pin1,3;4;7* and *pin1,3;4;7* was significant at *P*<0.001 (***) by Kruskal-Wallis and Mann-Whitney test with Bonferroni correction. Sample population sizes: WT, 48; *pin3;4;7*, 45; *pin1,3;4;7*, 70; PIN1::gPIN1:GFP;*pin3;4;7*, 60; ATML1::cPIN1:GFP;*pin3;4;7*, 37; PIN1::cPIN1:GFP;*pin3;4;7*, 28; SHR::cPIN1:GFP;*pin3;4;7*, 50; SCL32::cPIN1:GFP;*pin3;4;7*, 38; PIN1::gPIN1:GFP;*pin1,3;4;7*, 45; ATML1::cPIN1:GFP;*pin1,3;4;7*, 57; PIN1::cPIN1:GFP;*pin1,3;4;7*, 53; SHR::cPIN1:GFP;*pin1,3;4;7*, 62; SCL32::cPIN1:GFP;*pin1,3;4;7*, 69. e, epidermis. Scale bars: (A–C,E–I,K,L) 60 μm; (D,J) 20 μm; (M,N,O) 0.75 mm.

As previously shown (Verna et al., 2019), the vein pattern of *pin3;4;7* was no different from that of WT, and none of the *pin1,3;4;7* leaves had a WT vein pattern (Fig. 4M–P). The vein patterns of PIN1::gPIN1:GFP;*pin3;4;7*, ATML1::cPIN1:GFP;*pin3;4;7*, PIN1::cPIN1:GFP;*pin3;4;7*, SHR::cPIN1:GFP;*pin3;4;7*, and SCL32::cPIN1:GFP;*pin3;4;7* were no different from the WT vein pattern (Fig. 4M–P). Both PIN1::gPIN1:GFP and PIN1::cPIN1:GFP normalized the phenotype spectrum of *pin1,3;4;7* vein patterns (Fig. 4M–P; Fig. S3A,C). SHR::cPIN1:GFP shifted the phenotype spectrum of *pin1,3;4;7* vein patterns toward the WT vein network pattern, to match the phenotype spectrum of *pin1* vein patterns (Fig. 4M–P; Fig. S3D; cf. Fig. 2M–P). The vein pattern defects of ATML1::cPIN1:GFP;*pin1,3;4;7* and SCL32::cPIN1:GFP;*pin3;4;7* were no different from those of *pin1,3;4;7* (Fig. 4M– P; Fig. S3B,E). We observed a similar effect of tissue-specific *PIN1* expression on that component of cotyledon patterning that depends on *PIN1, PIN3, PIN4*, and *PIN7* (Figure S4).

Therefore, that PIN1 expression in the epidermis is dispensable and that PIN1 expression in the inner tissues of the leaf is sufficient for *PIN1*-dependent vein patterning cannot be accounted for by compensatory functions provided by *PIN3, PIN4*, and *PIN7*. Such compensatory functions are also unlikely provided by the remaining PIN proteins, by the ABCB1 and ABCB19 auxin efflux carriers, or by the AUX1/LAX auxin influx carriers because none of these proteins are either expressed in the epidermis or have functions in vein patterning, whether in WT or in auxin-transport-inhibited leaves (Verna et al., 2015; Sawchuk et al., 2013; Verna et al., 2019). As such, we conclude that PIN1 expression in the epidermis is dispensable for auxin-transport-dependent vein patterning. This conclusion is consistent with the observation that *cup-shaped cotyledon2* mutants lack convergent points of epidermal PIN1 polarity and yet have normal vein patterns (Bilsborough et al., 2011). By contrast, PIN1 expression in inner tissues is required and sufficient for auxin-transport-dependent vein patterning; such function of PIN1 expression seems to mainly depend on PIN1 expression in the vascular tissue.

In conclusion, vein patterning hypotheses based on polar auxin transport from the epidermis (reviewed in (Runions, Smith, & Prusinkiewicz, 2014; Prusinkiewicz & Runions, 2012); see also (Alim & Frey, 2010; Hartmann et al., 2019), and references therein) are unsupported by experimental evidence. Our results do not rule out an influence of the epidermis on vein patterning, for example through local auxin production (e.g., (Abley, Sauret-Gueto, Maree, & Coen, 2016)), but they do exclude that such influence is brought about by polar auxin transport. Alternatively, patterning of local epidermal features, such as peaks of auxin production or response, and of the processes that depend on those features may be mediated by auxin transport in underlying tissues; there is evidence for such possibility (e.g., (Deb, Marti, Frenz, Kuhlemeier, & Reinhardt, 2015)), and our results are consistent with that evidence. In the future, it will be interesting to test these and other possibilities, but already now our results refute all the vein patterning hypotheses that depend on polar auxin transport from the epidermis.

## Materials & Methods

### Notation

In agreement with (Crittenden, Bitgood, Burt, DW, Ponce de Leon, & Tixier-Boichard, 1996), linked genes or mutations (<2,500 kb apart, which in Arabidopsis on an average corresponds to ∼10 cM (Lukowitz, Gillmor, & Scheible, 2000)) are separated by a comma, and unlinked genes or mutations are separated by a semicolon.

### Plants

Origin and nature of lines, genotyping strategies, and oligonucleotide sequences are in Tables S1, S2, and S3, respectively. Seeds were sterilized and sown as in (Sawchuk, Donner, Head, & Scarpella, 2008). Stratified seeds were germinated and seedlings were grown at 22°C under continuous fluorescent light (∼80 µmol m^−2^s^−1^). Plants were grown at 25°C under fluorescent light (∼100 μmol m-2s-1) in a 16-h-light/8-h-dark cycle. Plants were transformed and representative lines were selected as in (Sawchuk et al., 2008).

### Imaging

Developing leaves were mounted and YFP was imaged as in (Sawchuk et al., 2013). CFP, YFP, and autofluorescence were imaged as in (Sawchuk et al., 2013). GFP and autofluorescence were imaged as in (Amalraj et al., 2019). Images were stacked, aligned with the Scale Invariant Feature Transform algorithm (Lowe, 2004), and maximum-intensity projection was applied to aligned image stacks in the Fiji distribution (Schindelin et al., 2012) of ImageJ (Schneider, Rasband, & Eliceiri, 2012; Schindelin, Rueden, Hiner, & Eliceiri, 2015; Rueden et al., 2017). Mature leaves were fixed in ethanol: acetic acid 6: 1, rehydrated in 70% ethanol and water, and mounted in chloral hydrate: glycerol: water 8: 2: 1. Mounted leaves were imaged as in (Odat et al., 2014). Greyscaled RGB color images were turned into 8-bit images, and image brightness and contrast were adjusted by linear stretching of the histogram in the Fiji distribution of ImageJ.

## Acknowledgements

We thank the Arabidopsis Biological Resource Center for PIN1::gPIN1:CFP, *PIN1* cDNA, and *pin1-1*; the Nottingham Arabidopsis Stock Centre for *pin1-051*; Ikram Blilou and Ben Scheres for *pin3-3, pin4-2*, and *pinγ^En^*; Megan Sawchuk for PIN1::nYFP; and Jian Xu and Ben Scheres for PIN1::gPIN1:YFP and PIN1::gPIN1:GFP. This work was supported by Discovery Grants of the Natural Sciences and Engineering Research Council of Canada (NSERC) to E.S. C.V. was supported, in part, by a University of Alberta Doctoral Recruitment Scholarship.

## Supplemental Figure Legends

**Figure S1. Effect of Tissue-Specific PIN1 Expression on *pin1* Vein Patterns.** (A-E) Dark-field illumination of mature first leaves. Scale bars: (A-E) 2 mm.

**Figure S2. Tissue-Specific PIN1 Expression in *PIN1*-dependent Cotyledon Patterning.** (A–G) Dark-field illumination of 3-day-old seedlings illustrating phenotype classes (top right): class I, two separate cotyledons (A); class II, two fused cotyledons and separate single cotyledon (B); class III, three fused cotyledons (C); class IV, three separate cotyledons (D); class V, two fused cotyledons (E); class VI, single cotyledon (F); class VII, cup-shaped cotyledon, side view (G). (H) Percentages of cotyledons in phenotype classes. Difference between *pin1* and WT, between PIN1::gPIN1:GFP;*pin1* and *pin1*, and between PIN1::cPIN1:GFP;*pin1* and *pin1* was significant at *P*<0.001 (***) by Kruskal-Wallis and Mann-Whitney test with Bonferroni correction. Sample population sizes: WT, 99; *pin1*, 50; PIN1::gPIN1:GFP, 110; ATML1::cPIN1:GFP, 113; PIN1::cPIN1:GFP, 115; SHR::cPIN1:GFP, 63; SCL32::cPIN1:GFP, 103; PIN1::gPIN1:GFP;*pin1*, 111; ATML1::cPIN1:GFP;*pin1*, 183; PIN1::cPIN1:GFP;*pin1*, 47; SHR::cPIN1:GFP;*pin1*, 45; SCL32::cPIN1:GFP;*pin1*, 54. Scale bars: (A–G) 0.5 mm.

**Figure S3. Effect of Tissue-Specific PIN1 Expression on *pin1,3;4;7* Vein Patterns.** (A-E) Dark-field illumination of mature first leaves. Scale bars: (A,C,D) 2 mm; (B,E) 1 mm.

**Figure S4. Tissue-Specific PIN1 Expression in *PIN1/PIN3/PIN4/PIN7*-dependent Cotyledon Patterning.** (A) Dark-field illumination of 3-day-old seedlings illustrating phenotype class VIII (top right) — small, hood-like outgrowth (side view). (H) Percentages of cotyledons in phenotype classes (classes I–VII defined in Figure S1). Difference between *pin1,3;4;7* and WT, between PIN1::gPIN1:PIN1;*pin1,3;4;7* and *pin1,3;4;7*, and between PIN1::cPIN1:PIN1;*pin1,3;4;7* and *pin1,3;4;7* was significant at *P*<0.001 (***) by Kruskal-Wallis and Mann-Whitney test with Bonferroni correction. Sample population sizes: WT, 102; *pin3;4;7*, 51; *pin1,3;4;7*, 130; PIN1::gPIN1:GFP;*pin3;4;7*, 65; ATML1::cPIN1:GFP;*pin3;4;7*, 108; PIN1::cPIN1:GFP;*pin3;4;7*, 107; SHR::cPIN1:GFP;*pin3;4;7*, 71; SCL32::cPIN1:GFP;*pin3;4;7*, 49; PIN1::gPIN1:GFP;*pin1,3;4;7*, 42; ATML1::cPIN1:GFP;*pin1,3;4;7*, 83; PIN1::cPIN1:GFP;*pin1,3;4;7*, 85; SHR::cPIN1:GFP;*pin1,3;4;7*, 60; SCL32::cPIN1:GFP;*pin1,3;4;7*, 49. Scale bar: (A) 0.25 mm.

## References

Abley, K., Sauret-Gueto, S., Maree, A. F., & Coen, E. (2016). Formation of polarity convergences underlying shoot outgrowths. Elife, 5, e18165.

Alim, K., & Frey, E. (2010). Quantitative predictions on auxin-induced polar distribution of PIN proteins during vein formation in leaves. The European Physical Journal E, 33(2), 165–173.

Amalraj, B., Govindaraju, P., Krishna, A., Lavania, D., Linh, N. M., Ravichandran, S. J. et al. (2019). GAL4/GFP enhancer-trap lines for identification and manipulation of cells and tissues in developing Arabidopsis leaves. bioRxiv, 801357.

Bayer, E. M., Smith, R. S., Mandel, T., Nakayama, N., Sauer, M., Prusinkiewicz, P. et al. (2009). Integration of transport-based models for phyllotaxis and midvein formation. Genes Dev, 23(3), 373–384.

Benkova, E., Michniewicz, M., Sauer, M., Teichmann, T., Seifertova, D., Jurgens, G. et al. (2003). Local, efflux-dependent auxin gradients as a common module for plant organ formation. Cell, 115(5), 591–602.

Bilsborough, G. D., Runions, A., Barkoulas, M., Jenkins, H. W., Hasson, A., Galinha, C. et al. (2011). Model for the regulation of Arabidopsis thaliana leaf margin development. Proc Natl Acad Sci U S A, 108(8), 3424–3429.

Cherbas, L., Hu, X., Zhimulev, I., Belyaeva, E., & Cherbas, P. (2003). EcR isoforms in Drosophila: testing tissue-specific requirements by targeted blockade and rescue. Development, 130(2), 271–284.

Crittenden, L. B., Bitgood, J. J., Burt, DW, Ponce de Leon, F. A., & Tixier-Boichard, M. (1996). Nomenclature for naming loci, alleles, linkage groups and chromosomes to be used in poultry genome publications and databases. Genet Sel Evol, 28, 289–297.

Deb, Y., Marti, D., Frenz, M., Kuhlemeier, C., & Reinhardt, D. (2015). Phyllotaxis involves auxin drainage through leaf primordia. Development, 142(11), 1992–2001.

Donner, T. J., & Scarpella, E. (2013). Transcriptional control of early vein expression of CYCA2; 1 and CYCA2;4 in Arabidopsis leaves. Mech Dev, 130(1), 14–24.

Donner, T. J., Sherr, I., & Scarpella, E. (2009). Regulation of preprocambial cell state acquisition by auxin signaling in Arabidopsis leaves. Development, 136(19), 3235–3246.

Galweiler, L., Guan, C., Muller, A., Wisman, E., Mendgen, K., Yephremov, A. et al. (1998). Regulation of polar auxin transport by AtPIN1 in Arabidopsis vascular tissue. Science, 282(5397), 2226–2230.

Gardiner, J., Donner, T. J., & Scarpella, E. (2011). Simultaneous activation of SHR and ATHB8 expression defines switch to preprocambial cell state in Arabidopsis leaf development. Dev Dyn, 240(1), 261–270.

Gardiner, J., Sherr, I., & Scarpella, E. (2010). Expression of DOF genes identifies early stages of vascular development in Arabidopsis leaves. Int J Dev Biol, 54(8-9), 1389–1396.

Gordon, S. P., Heisler, M. G., Reddy, G. V., Ohno, C., Das, P., & Meyerowitz, E. M. (2007). Pattern formation during de novo assembly of the Arabidopsis shoot meristem. Development, 134(19), 3539–3548.

Hartmann, F. P., Barbier de Reuille, P., & Kuhlemeier, C. (2019). Toward a 3D model of phyllotaxis based on a biochemically plausible auxin-transport mechanism. PloS Comp Biol, 15(4), e1006896.

Hay, A., Barkoulas, M., & Tsiantis, M. (2006). ASYMMETRIC LEAVES1 and auxin activities converge to repress BREVIPEDICELLUS expression and promote leaf development in Arabidopsis. Development, 133(20), 3955–3961.

Heisler, M. G., Ohno, C., Das, P., Sieber, P., Reddy, G. V., Long, J. A. et al. (2005). Patterns of Auxin Transport and Gene Expression during Primordium Development Revealed by Live Imaging of the Arabidopsis Inflorescence Meristem. Curr Biol, 15(21), 1899–1911.

Kang, J., & Dengler, N. (2004). Vein pattern development in adult leaves of Arabidopsis thaliana. International Journal of Plant Sciences, 165(2), 231–242.

Lowe, D. G. (2004). Distinctive image features from scale-invariant keypoints. International journal of computer vision, 60(2), 91–110.

Lukowitz, W., Gillmor, C. S., & Scheible, W. R. (2000). Positional cloning in Arabidopsis. Why it feels good to have a genome initiative working for you. Plant Physiol, 123(3), 795–805.

Marcos, D., & Berleth, T. (2014). Dynamic auxin transport patterns preceding vein formation revealed by live-imaging of Arabidopsis leaf primordia. Front Plant Sci, 5, 235.

Mattsson, J., Sung, Z. R., & Berleth, T. (1999). Responses of plant vascular systems to auxin transport inhibition. Development, 126(13), 2979–2991.

Noden, D. M. (1988). Interactions and fates of avian craniofacial mesenchyme. Development, 103 Supplement, 121–140.

Odat, O., Gardiner, J., Sawchuk, M. G., Verna, C., Donner, T. J., & Scarpella, E. (2014). Characterization of an allelic series in the MONOPTEROS gene of Arabidopsis. Genesis, 52(2), 127–133.

Petrasek, J., Mravec, J., Bouchard, R., Blakeslee, J. J., Abas, M., Seifertova, D. et al. (2006). PIN proteins perform a rate-limiting function in cellular auxin efflux. Science, 312(5775), 914–918.

Prusinkiewicz, P., & Runions, A. (2012). Computational models of plant development and form. New Phytol, 193(3), 549–569.

Reinhardt, D., Pesce, E. R., Stieger, P., Mandel, T., Baltensperger, K., Bennett, M. et al. (2003). Regulation of phyllotaxis by polar auxin transport. Nature, 426(6964), 255–260.

Rueden, C. T., Schindelin, J., Hiner, M. C., DeZonia, B. E., Walter, A. E., Arena, E. T. et al. (2017). ImageJ2: ImageJ for the next generation of scientific image data. BMC Bioinformatics, 18(1), 529.

Runions, A., Smith, R. S., & Prusinkiewicz, P. (2014). Computational Models of Auxin-Driven Development. In E. Zažímalová, J. Petrasek, & E. Benková (Eds.), Auxin and Its Role in Plant Development (pp. 315–357).

Sawchuk, M. G., Donner, T. J., Head, P., & Scarpella, E. (2008). Unique and overlapping expression patterns among members of photosynthesis-associated nuclear gene families in Arabidopsis. Plant Physiol, 148(4), 1908–1924.

Sawchuk, M. G., Edgar, A., & Scarpella, E. (2013). Patterning of leaf vein networks by convergent auxin transport pathways. PLoS Genet, 9(2), e1003294.

Sawchuk, M. G., Head, P., Donner, T. J., & Scarpella, E. (2007). Time-lapse imaging of Arabidopsis leaf development shows dynamic patterns of procambium formation. New Phytol, 176(3), 560–571.

Scarpella, E., Marcos, D., Friml, J., & Berleth, T. (2006). Control of leaf vascular patterning by polar auxin transport. Genes Dev, 20(8), 1015–1027.

Scarpella, E., Francis, P., & Berleth, T. (2004). Stage-specific markers define early steps of procambium development in Arabidopsis leaves and correlate termination of vein formation with mesophyll differentiation. Development, 131(14), 3445–3455.

Schindelin, J., Arganda-Carreras, I., Frise, E., Kaynig, V., Longair, M., Pietzsch, T. et al. (2012). Fiji: an open-source platform for biological-image analysis. Nat Methods, 9(7), 676–682.

Schindelin, J., Rueden, C. T., Hiner, M. C., & Eliceiri, K. W. (2015). The ImageJ ecosystem: An open platform for biomedical image analysis. Mol Reprod Dev, 82(7-8), 518–529.

Schneider, C. A., Rasband, W. S., & Eliceiri, K. W. (2012). NIH Image to ImageJ: 25 years of image analysis. Nat Methods, 9(7), 671–675.

Sessions, A., Weigel, D., & Yanofsky, M. F. (1999). The Arabidopsis thaliana MERISTEM LAYER 1 promoter specifies epidermal expression in meristems and young primordia. Plant J, 20(2), 259–263.

Sieburth, L. E. (1999). Auxin is required for leaf vein pattern in Arabidopsis. Plant Physiol, 121(4), 1179–1190.

Soloviev, A., Gallagher, J., Marnef, A., & Kuwabara, P. E. (2011). C. elegans patched-3 is an essential gene implicated in osmoregulation and requiring an intact permease transporter domain. Dev Biol, 351(2), 242–253.

Topalidou, I., & Miller, D. L. (2017). <i>Caenorhabditis elegans</i> HIF-1 Is Broadly Required for Survival in Hydrogen Sulfide. G3 (Bethesda), 7(11), 3699–3704.

Verna, C., Sawchuk, M. G., Linh, N. M., & Scarpella, E. (2015). Control of vein network topology by auxin transport. BMC Biol, 13, 94.

Verna, C., Ravichandran, S. J., Sawchuk, M. G., Linh, N. M., & Scarpella, E. (2019). Coordination of Tissue Cell Polarity by Auxin Transport and Signaling. Elife, 8, e51061.

Wenzel, C. L., Schuetz, M., Yu, Q., & Mattsson, J. (2007). Dynamics of MONOPTEROS and PIN-FORMED1 expression during leaf vein pattern formation in Arabidopsis thaliana. Plant J, 49(3), 387–398.

Wisidagama, D. R., Thomas, S. M., Lam, G., & Thummel, C. S. (2019). Functional analysis of Aarf domain-containing kinase 1 in Drosophila melanogaster. Dev Dyn, 248(9), 762–770.

Xu, J., Hofhuis, H., Heidstra, R., Sauer, M., Friml, J., & Scheres, B. (2006). A molecular framework for plant regeneration. Science, 311(5759), 385–388.

Xue, Y., Gao, X., Lindsell, C. E., Norton, C. R., Chang, B., Hicks, C. et al. (1999). Embryonic lethality and vascular defects in mice lacking the Notch ligand Jagged1. Hum Mol Genet, 8(5), 723–730.

Zourelidou, M., Absmanner, B., Weller, B., Barbosa, I. C., Willige, B. C., Fastner, A. et al. (2014). Auxin efflux by PIN-FORMED proteins is activated by two different protein kinases, D6 PROTEIN KINASE and PINOID. Elife, 3, e02860.

