## Supplementary material for "Vein Patterning by Tissue-Specific Auxin Transport": Table S1

**Table 1. Origin and Nature of Lines.**

| Line | Origin/Nature |
| --- | --- |
| PIN1::gPIN1:YFP | (Xu et al., 2006) |
| PIN1::nYFP | Transcriptional fusion of *PIN1* (AT1G73590; -4,171 to -1; primers: "PIN1 transc 4171 forw" and "PIN1 transc rev") to HTA6:EYFP (Zhang et al., 2005) |
| PIN1::gPIN1:CFP | (Gordon et al., 2007) |
| *pin1-051* | NASC; GK-051A10-012139 (Kleinboelting et al., 2012); contains a T-DNA insertion after +2234 of *PIN1* |
| PIN1::gPIN1:GFP | Xu et al. 2006 |
| ATML1::cPIN1:GFP | Transcriptional fusion of *ATML1* (AT4G21750; -5,016 to -1,597; primers "XhoI ATML1 p F" and "BamHI ATML1p R") to translational fusion of *PIN1* cDNA (GenBank accession no. AY093960; ABRC clone no. U12338; primers "BamHI PIN1 cDNA F" and "KpnI PIN1 cDNA R") to EGFP (Clontech; insertion after +651 of *PIN1*; primers "XhoI GFP no ATG Fwd" and "XhoI GFP no* Rev") |
| PIN1::cPIN1:GFP | Transcriptional fusion of *PIN1* (-4,168 to -14; primers "XhoI full length PIN1p F" and "BamHI PIN1p rev") to translational fusion of *PIN1* cDNA (GenBank accession no. AY093960; ABRC clone no. U12338; primers "BamHI PIN1 cDNA F" and "KpnI PIN1 cDNA R") to EGFP (Clontech; insertion after +651 of *PIN1*; primers "XhoI GFP no ATG Fwd" and "XhoI GFP no* Rev") |

| | |
|---|---|
| SHR::cPIN1:GFP | Transcriptional fusion of *SHR* (AT4G37650; -2505 to -16; primers "SHR prom SalI Forw2" and "SHR prom BamHI Rev") to translational fusion of *PIN1* cDNA (GenBank accession no. AY093960; ABRC clone no. U12338; primers "BamHI PIN1 cDNA F" and "KpnI PIN1 cDNA R") to EGFP (Clontech; insertion after +651 of *PIN1*; primers "XhoI GFP no ATG Fwd" and "XhoI GFP no* Rev") |
| SCL32::cPIN1:GFP | Transcriptional fusion of *SCL32* (AT3G49950; -2888 to -2; primers "SCL32 Translational FWD" and "SCL32 prom BamHI Rev") to translational fusion of *PIN1* cDNA (GenBank accession no. AY093960; ABRC clone no. U12338; primers "BamHI PIN1 cDNA F" and "KpnI PIN1 cDNA R") to EGFP (Clontech; insertion after +651 of *PIN1*; primers "XhoI GFP no ATG Fwd" and "XhoI GFP no* Rev") |
| PIN3::gPIN3:YFP | ABRC; (Zhou et al., 2011) |
| PIN4::gPIN4:YFP | ABRC; (Zhou et al., 2011) |
| PIN7::gPIN7:YFP | ABRC; (Zhou et al., 2011) |
| *pin1-1* | ABRC; WT at the *TTG1* (AT5G24520) locus (Galweiler et al., 1998; Goto N, 1987; Sawchuk et al., 2013) |
| *pin3-3* | (Friml et al., 2002b) |
| *pin4-2* | (Friml et al., 2002a) |
| $pin7^{En}$ | (Blilou et al., 2005) |

Blilou, I., Xu, J., Wildwater, M., Willemsen, V., Paponov, I., Friml, J., Heidstra, R., Aida, M., Palme, K., Scheres, B., 2005. The PIN auxin efflux facilitator network controls growth and patterning in Arabidopsis roots. Nature 433, 39-44.

Friml, J., Benkova, E., Blilou, I., Wisniewska, J., Hamann, T., Ljung, K., Woody, S., Sandberg, G., Scheres, B., Jurgens, G., Palme, K., 2002a. AtPIN4 mediates sink-driven auxin gradients and root patterning in Arabidopsis. Cell 108, 661-673.

Friml, J., Wisniewska, J., Benkova, E., Mendgen, K., Palme, K., 2002b. Lateral relocation of auxin efflux regulator PIN3 mediates tropism in Arabidopsis. Nature 415, 806-809.

Galweiler, L., Guan, C., Muller, A., Wisman, E., Mendgen, K., Yephremov, A., Palme, K., 1998. Regulation of polar auxin transport by AtPIN1 in Arabidopsis vascular tissue. Science 282, 2226-2230.

Gordon, S.P., Heisler, M.G., Reddy, G.V., Ohno, C., Das, P., Meyerowitz, E.M., 2007. Pattern formation during de novo assembly of the Arabidopsis shoot meristem. Development 134, 3539-3548.

Goto N, S.M., Kranz AR, 1987. Effect of gibberllins on flower development of the pin-formed mutant of Arabidopsis thaliana. Arabidopsis Information Service 23, 66-71.

Kleinboelting, N., Huep, G., Kloetgen, A., Viehoever, P., Weisshaar, B., 2012. GABI-Kat SimpleSearch: new features of the Arabidopsis thaliana T-DNA mutant database. Nucleic Acids Res 40, D1211-5.

Sawchuk, M.G., Edgar, A., Scarpella, E., 2013. Patterning of leaf vein networks by convergent auxin transport pathways. PLoS Genet 9, e1003294.

Xu, J., Hofhuis, H., Heidstra, R., Sauer, M., Friml, J., Scheres, B., 2006. A molecular framework for plant regeneration. Science 311, 385-388.

Zhang, C., Gong, F.C., Lambert, G.M., Galbraith, D.W., 2005. Cell type-specific characterization of nuclear DNA contents within complex tissues and organs. Plant Methods 1, 7.

Zhou, R., Benavente, L.M., Stepanova, A.N., Alonso, J.M., 2011. A recombineering-based gene tagging system for Arabidopsis. Plant J 66, 712-723.
