## Supplementary material for "Vein Patterning by Tissue-Specific Auxin Transport": Table S2

**Table 2. Genotyping Strategies.**

| Line | Strategy |
|---|---|
| *pin1-051* | *PIN1*: "pin1 GK LP" and "pin1 GK RP"; *pin1:* "pin1 GK RP" and "o8409" |
| *pin1-1* | "pin1-1 F" and "pin1-1 R"; *Tat*I |
| *pin3-3* | "pin3-3 F" and "pin3-3 R"; *Sty*I |
| *pin4-2* | *PIN4*: "PIN4 forw geno II" and "PIN4en rev Ikram"; *pin4*: "PIN4en rev Ikram" and "en primer" |
| *pin7*$^{En}$ | *PIN7*: "PIN7en forw Ikram" and "PIN7en rev"; *pin7*: "PIN7en rev Ikram II" and "en primer" |
