## Supplementary material for "Vein Patterning by Tissue-Specific Auxin Transport": Table S3

**Table 3. Oligonucleotide Sequences.**

| Name | Sequence (5' to 3') |
|---|---|
| PIN1 transc 4171 forw | GGGGACAAGTTTGTACAAAAAAGCAGGCTATGATC CGATTGGATTCG |
| PIN1 transc rev | GGGGACCACTTTGTACAAGAAAGCTGGGTCTTTTG TTCGCCGGAGAAG |
| pin GK LP | ACTCTTTGGCAAACACAAACG |
| pin1 GK RP | CTCTCAGATGCAGGTCTAGGC |
| o8409 | ATATTGACCATCATACTCATTGC |
| XhoI ATML1 p F | GCCCTCGAGTTTACATTGATTCTGAACTG |
| BamHI ATML1p R | GATGGATCCTAACCGGTGGATTCAGGGAG |
| BamHI PIN1 cDNA F new | TTAGGATCCATGATTACGGCGGCGGACTTC |
| KpnI PIN1 cDNA R | CTCGGTACCTCATAGACCCAAGAGAATGTAG |
| XhoI GFP no ATG Fwd | TTACTCGAGAGTGAGCAAGGGCGAGGAGCTGTT |
| XhoI GFP no* Rev | TATCTCGAGTACTTGTACAGCTCGTCCATGCCGAG |
| XhoI full length PIN1p F | TGTCTCGAGATCCGATTGGATTCGGTCTG |
| BamHI PIN1p rev | AAGGGATCCGAGAAGAGAGAGGGAAGAGAG |
| SHR prom SalI Forw2 | AAAGTCGACCGAAGAAAGGGACAAAGAAGC |
| SHR prom BamHI Rev | TGGGGATCCTTAATGAATAAGAAAATGAATAGAAG AAAGGG |
| SCL32 Translational FWD | AGAGTCGACATCTTAGTAGAAATAAGCGAAC |
| SCL32 prom BamHI Rev | ACTGGATCCGAGTCTGGTTTTAGAGAGAAATG |
| pin1-1 F | ATGATTACGGCGGCGGACTTCTA |
| pin1-1 R | TTCCGACCACCACCAGAAGCC |
| pin3-3 F | GGAGCTCAAACGGGTCACCCG |
| pin3-3 R | GCTGGATGAGCTACAGCTATATTC |
| PIN4 forw geno II | GTCCGACTCCACGGCCTTC |
| PIN4en rev Ikram | ATCTTCTTCTTCACCTTCCACTCT |
| en primer | GAGCGTCGGTCCCCACACTTCTATAC |
| PIN7en forw Ikram | CCTAACGGTTTCCACACTCA |
| PIN7en rev | TAGCTCTTTAGGGTTTAGCTC |
| PIN7en rev Ikram II | GGTTTAGCTCTGCTGTGGAGTT |
