## Supplementary figures and images for "Vein Patterning by Tissue-Specific Auxin Transport"

### Figure S1

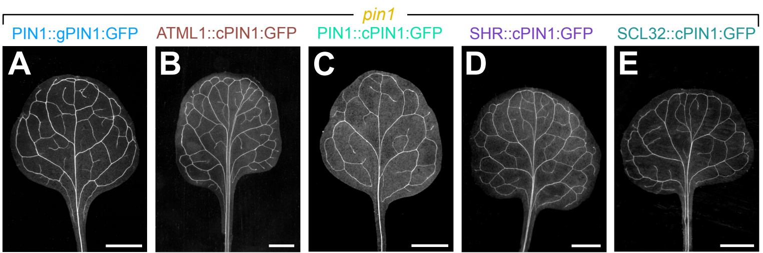

### Figure S2

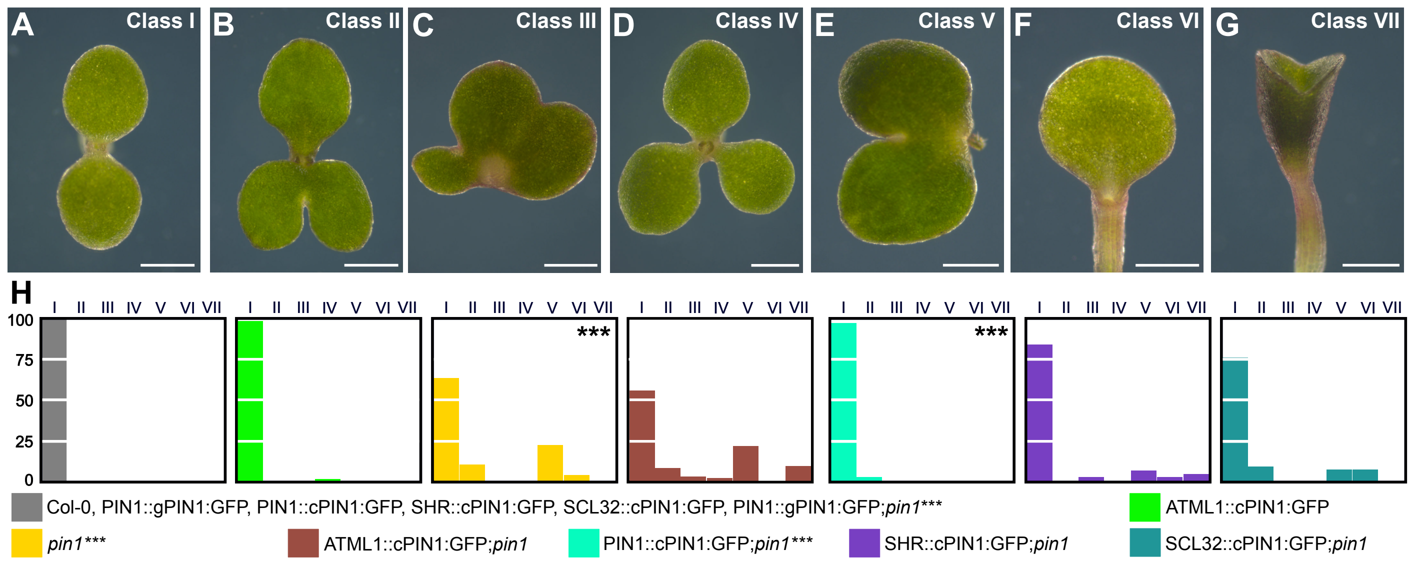

### Figure S3

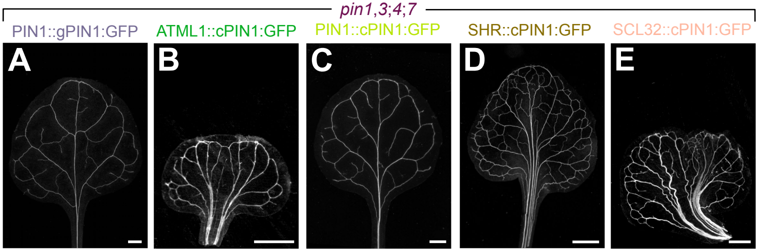

### Figure S4

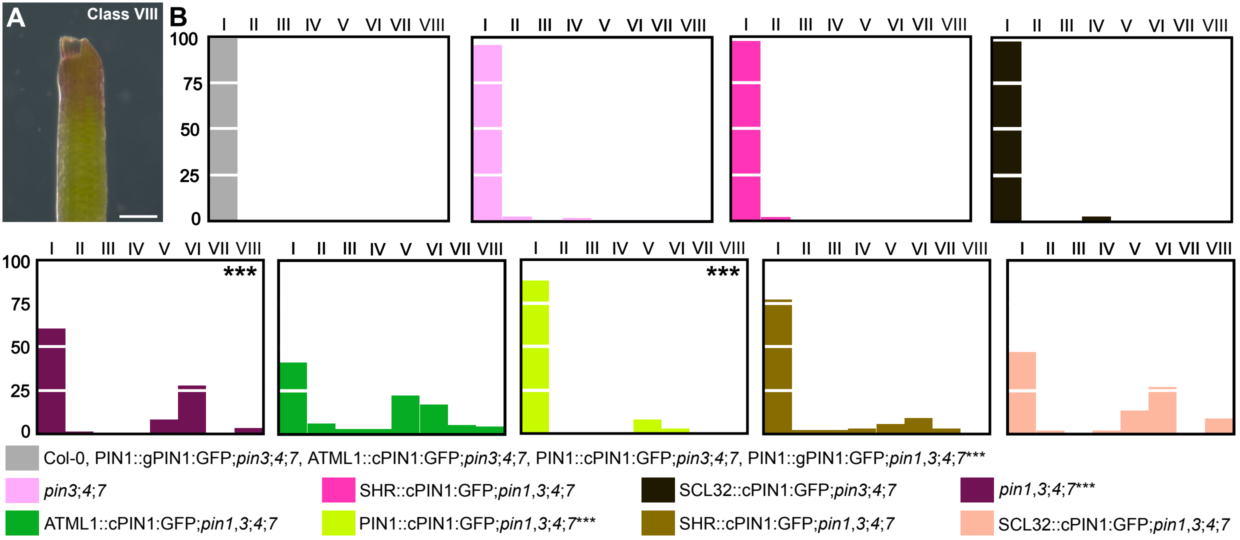
